# NERDSS: a non-equilibrium simulator for multibody self-assembly at the cellular scale

**DOI:** 10.1101/853614

**Authors:** Matthew J. Varga, Spencer Loggia, Yiben Fu, Osman N Yogurtcu, Margaret E. Johnson

## Abstract

Currently, a significant barrier to building predictive models of cell-based self-assembly processes is that molecular models cannot capture minutes-long cellular dynamics that couple distinct components with active processes, while reaction-diffusion models lack sufficient detail for capturing assembly structures. Here we introduce the Non-Equilibrium Reaction-Diffusion Self-assembly Simulator (NERDSS), which addresses this gap by integrating a structure-resolved reaction-diffusion algorithm with rule-based model construction. By representing proteins as rigid, multi-site molecules that adopt well-defined orientations upon binding, NERDSS simulates formation of large reversible structures with sites that can be acted on by reaction rules. We show how NERDSS allows for directly comparing and optimizing models of multi-component assembly against time-dependent experimental data. Applying NERDSS to assembly steps in clathrin-mediated endocytosis, we capture how the formation of clathrin caged structures can be driven by modulating the strength of clathrin-clathrin interactions, by adding cooperativity, or by localizing clathrin to the membrane. NERDSS further predicts how clathrin lattice disassembly can be driven by enzymes that irreversibly change lipid populations on the membrane. By modeling viral lattice assembly and recapitulating oscillations in protein expression levels for a circadian clock model, we illustrate the wide usability and adaptability of NERDSS. NERDSS simulates user-defined assembly models that were previously inaccessible to existing software tools, with broad applications to predicting self-assembly *in vivo* and designing high-yield assemblies *in vitro*.

## Introduction

Watching the dynamics of individual molecules in the cell as they function as part of a collective is now possible thanks to revolutions in live-cell microscopy. Computer simulations offer the promise of reproducing these dynamics using models designed from the underlying physics and mechanics, providing high resolution spatial and temporal predictions and exquisite control over components and their interactions. Current tools for capturing cell-scale complexity are a growing companion to cell biology. For example, the field of cell-signaling has benefitted from a variety of spatial^1, 2^ and non-spatial tools^3, 4^ where interactions and reactions are modeled as events parameterized by rate constants. However, many cell-scale processes involve self-assembly—a challenge for modeling because it spans similarly long length and time-scales as signaling cascades while also depending fundamentally on molecular structural geometry. Rate-based tools have been applied to self-assembly, but lack spatial resolution^3, 5^, apply only to very small systems^6, 7^, or include potentials that prevent quantitative comparison to experiment^8^. Here, we present the Non-Equilibrium Reaction-Diffusion Self-assembly Simulator (NERDSS), a higher-resolution rate-based software tool with addition of user-specified, coarse molecular structure to enable the study of self-assembly at the cell scale.

NERDSS was developed to build self-assembly into the reaction-diffusion (RD) model, because of the strength of RD in simulating relatively slow, non-equilibrium cellular dynamics. The length and time-scales of such dynamics are difficult or impossible with the alternative and standard computational approach for self-assembly of coarse-grained molecular dynamics (MD)-type models^9–11^. For those models, interactions emerge due to distance-dependent energy functions rather than rate-controlled events, so while they are more physically realistic, they lack systematic and transferable methods for involving enzymatic or ATP-driven reactions ubiquitous in cells. They are unable to simulate *in vivo* cell-signaling, cytoskeletal dynamics, or clathrin-mediated endocytosis, for example. Our NERDSS software addresses this substantial application gap, and while it lacks the detail of energy function based models, it is able to uniquely preserve important features of molecular assembly.

NERDSS overcomes several technical challenges with adding structure to RD and allowing large, reversible complexes to form, in a user-friendly and extensible way. To extend beyond single-particle RD^12–15^, proteins here can have multiple interaction sites at specific coordinates, and to reach large system sizes, we use the Free-Propagator Reweighting algorithm^12^, with recent extensions to include and account for rotational as well as translation diffusion^16^. To maintain accurate solutions to the RD equations of motion, and direct comparison to experimental rates, binding association between proteins can occur upon collision and does not depend on their relative orientations^16^. Bound proteins are instead oriented according to user-defined geometries once association events occur, which can be designed by our GUI. This also retains the flexibility and the adaptability of the method to new molecules and structures, as binding is parameterized by rates and not interaction potentials. NERDSS preserves detailed-balance for binding and unbinding between discrete species, or between proteins co-localized through a series of shared binding partners, thus allowing thermodynamic equilibrium for reversible systems, both on and off the membrane^17, 18^. Even large assemblies are thus capable of coming apart spontaneously, or with NERDSS dissociation can be catalyzed by additional reactions. The rule-based^19^ implementation of reactions means that users can choose whether to add or delete cooperativity, enzymatic reactions, or conditionality to binding interactions, providing a tool to probe a broad range of assembly mechanisms.

We apply NERDSS here to modeling virion assembly, protein expression dynamics, and clathrin-mediated endocytosis (CME), with a special emphasis on the latter. These examples highlight several challenges and opportunities for modeling approaches. CME is an essential pathway used by all eukaryotes for transport across the plasma membrane. A wealth of biochemical^20^, structural^21^, and *in vivo* imaging data ^22–24^ is thus available, but predicting cargo uptake by CME remains remarkably difficult because of the complexity of the pathway. CME is a stochastic process that depends on membrane mechanics, enzymatic reactions, and the stoichiometry of dozens of distinct components. Clathrin, a 600kDa trimeric protein, assembles into both flat and spherical lattices *in vivo* and *in vitro*, although *in vivo* this process only occurs on the plasma membrane, with the requirement of a host of accessory and adaptor proteins, as clathrin itself does not bind the membrane. Existing simulations of clathrin-cage assembly provide important physical insight into CME, but they use ad hoc energy-based models that require substantial expertise to develop^11,25, 26^, or lack spatial resolution^27, 28^. Here we implement an extensible multi-component model of clathrin lattice assembly to quantify distinct pathways and dynamics of assembly in solution and on the membrane, and directly reproduce fluorescence experiments^29^. Our simulations also show how potently the membrane can act to regulate assembly simply through reduction of dimensionality (3D to 2D)^18^. With NERDSS, we are able to predict how the dephosphorylation of essential plasma membrane lipids, which occurs at sites of vesicle budding^22^, can de-stabilize clathrin-coated structures and drive their disassembly.

In summary, NERDSS offers a distinct tool that uses the reaction-diffusion model to simulate self-assembly at the cell scale, thus allowing for space and time-dependent dynamics and kinetics that are immediately comparable to experiment by a broad user base. Below, we first describe the operation of the software and the features we introduced as necessary to move beyond existing tools (Fig 1). We then present multiple applications, with input files provided in the software repository. We develop several models of clathrin lattice assembly, highlighting distinct features of the models that can be used to tune the dynamics and stability of the observed structures. We optimize a clathrin model directly against time-dependent *in vitro* experimental data of clathrin recruitment and assembly on membranes^29^. We further introduce enzymes to the system to control the lipid populations on the membrane and drive disassembly, capturing driven nonequilibrium dynamics. We illustrate the model design process for the self-assembly of Gag retroviral protein monomers into an immature lattice, which is an essential component of the HIV infection and maturation cycle^30^. Our addition here of self-assembly to reaction-diffusion models augments the existing range of RD simulation capabilities. To demonstrate use of NERDSS for non-assembly problems, we recapitulate the oscillations of protein expression in a circadian clock model^31^, with validation against simulations using Virtual Cell software^1^. Finally, we discuss the current limitations and most promising future advancements of NERDSS software for realistic cell-scale dynamics.

**Figure 1:**
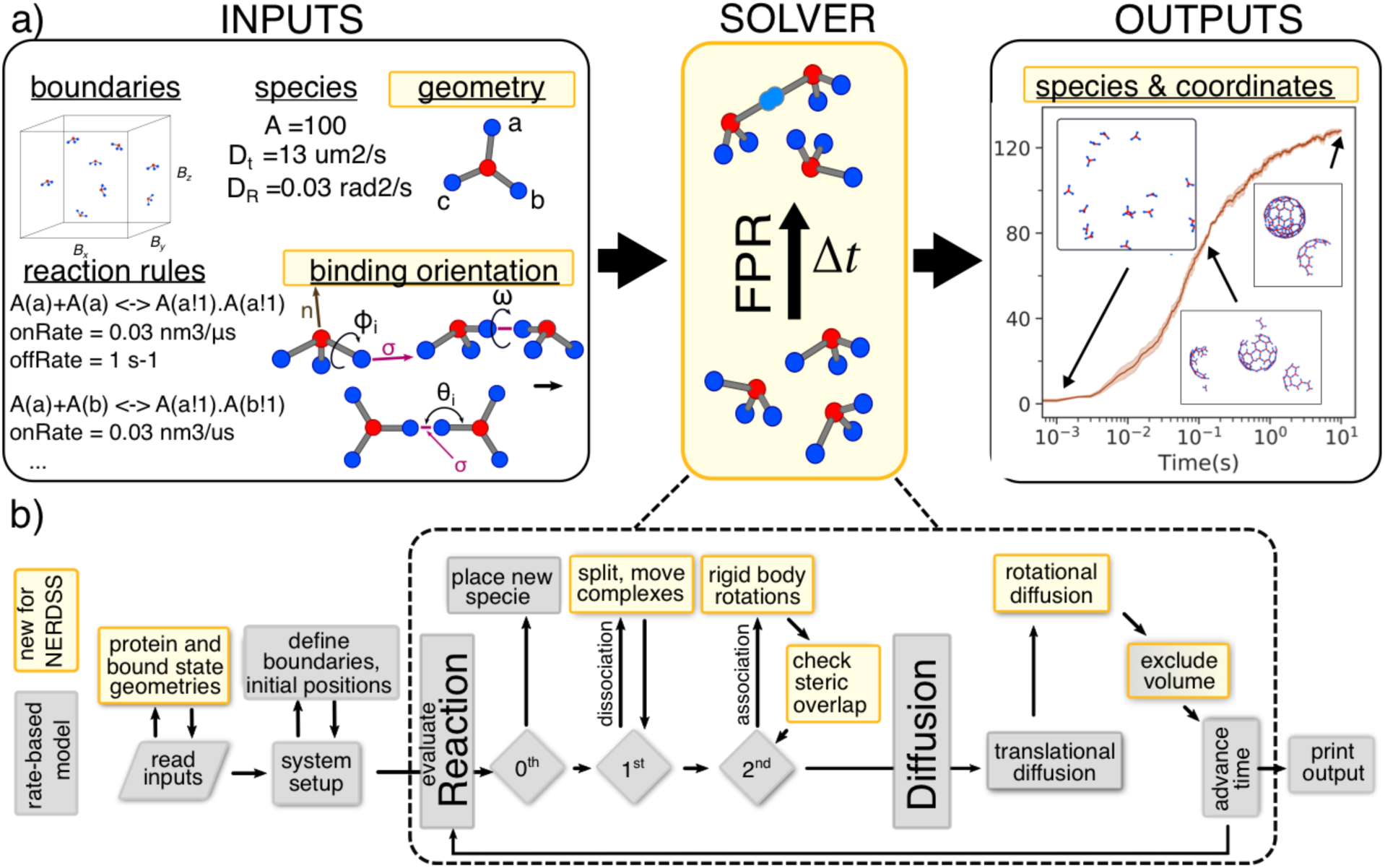
NERDSS software overview. a) NERDSS requires several familiar user-defined inputs, including simulation boundary conditions, species definitions, and reaction rules for these species written in BNGL-style syntax^19^ to act on individual interfaces or on entire molecules. For NERDSS, additional inputs in yellow boxes are the species geometry, with each specie modeled as a rigid body with coordinates for a center of mass (red) and its discrete interfaces (blue), and the binding orientation. These orientation parameters define the geometry of the resulting complex: a binding radius, σ, shown as a purple line between two interfaces; two angles, *θ*_1_ and *θ*_2_; three dihedral angles, *φ*_1_, *φ*_2_, and *ω* (see Supporting Information). Both geometry and orientation can be designed with the NERDSS GUI. NERDSS uses free-propagator reweighting algorithms (FPR) to solve the reaction-diffusion model over time. In addition to coordinates of all species in the simulation as a function of time, NERDSS can track a variety of variables, including species copy numbers (or current number of interactions) which are written to CSV formatted files for analysis and visualization. Trajectories are output in one of two standard formats, XYZ or PDB, which can be visualized in software such as VMD^32^ or Ovito^33^. b) Algorithmic flow-chart highlights where modifications to time and/or space dependent rate-based models (grey) are needed due to addition of molecular structure (yellow). Excluded volume is enforced in a small subset of spatial methods, hence the dual color. Reactions can be 0^th^ order: creation, 1^st^ order: death, transformations, dissociations, 2^nd^ order: binding, to a bound state or to two products. For each time-step, every molecule either reacts or diffuses.

## Results

### NERDSS enables self-assembly simulations

NERDSS combines recent algorithms for structure-resolved reaction diffusion^16^ with rule-based modeling^19^, orientable components, and user-friendly tools for system design (Fig 1a), to create broadly generalizable reaction-diffusion software capable of simulating self-assembly. NERDSS is efficient enough to simulate seconds to minutes of assembly on a single processer (Fig S1). Validation of NERDSS is done via changing time-steps^16^ (Fig S2) and by comparison of models against theory and other rate-based methods (Fig S3-S5). Several features of NERDSS are new for single-particle and spatial RD (Fig 1b), including representing molecules (such as proteins or lipids) as discrete objects with interfaces positioned at specific coordinates relative to a molecule center-of-mass.

Including finer geometry of molecular contacts into self-assembly creates two challenges. First, simulation algorithms must correctly simulate rotation, diffusion, and the reaction into the proper geometry. In NERDSS, molecules can bind one another upon collision, similar to other single-particle RD methods^12–14^, but because they can have a 3D structure, association is accompanied by orientation of the molecules into a bound state. Our recently developed algorithm propagates RD dynamics that includes both rotational and translational diffusion^16^. A unique feature of NERDSS is steric overlap evaluation upon association. Although volume exclusion is enforced during propagation between all interface pairs that react (which is shared with some other single-particle algorithms^12, 13^), association events can result in two multi-component assemblies being snapped into a bound state that forces interfaces to overlap with one another. These moves are rejected, as are moves that create assemblies too large to fit within the simulation volume. NERDSS allows binding between molecules regardless of their orientation prior to the event, and thus some reactions can produce large rotations for extended assemblies: restrictions can also be placed to reject these moves. Second, the user must be able to conveniently specify the geometry of the complex itself. The bound states are defined as part of the model during the system set-up. For example, this involves placing binding interfaces at a site-site separation (σ), and allowing for rotation of the two molecules by up to 5 relative angles into the designed orientation. We provide a GUI to facilitate the otherwise cumbersome construction of these molecules and bound states.

It is information to compare NERDSS to NFSim^3^, which lacks spatial resolution, but simulates rate-based models that can form multi-protein assemblies via multi-site species and rule-based modeling (Fig S3-S4). Similar to NFSim and other non-spatial rate-based methods, NERDSS uses rule-based modeling to specify reactions involving whole species, specific interfaces, or specific states of interfaces (e.g. phosphorylated or unphosphorylated). Rule-based modeling precludes the need to define and track all possible species and avoids the issue of combinatorial complexity^34^. The rule-based format translates quite naturally to NERDSS, due to the spatial specification of each interface, which is not true of other single-particle RD methods. With structural resolution, NERDSS captures several elements of self-assembly not possible with NFsim, including lattice geometries and topologies, and their growth patterns.

Distinct from non-spatial and spatial methods, NERDSS identifies when binding interfaces are localized within the same complex (as occurs e.g. in clathrin lattice formation) and uses spatial constraints to evaluate whether binding can occur. These intracomplex loop-closure reactions are effectively unimolecular, as the components are no longer diffusing in space relative to one another, and the rates are defined using a formula to preserve free energy barriers of pairwise component rates (Methods). With its spatial resolution, NERDSS also naturally adjusts for changes to reaction rates due to restriction of one or both components to the 2D surface (Methods). These reactions are solved accurately using FPR algorithms^12, 16, 17^ and are designed to preserve detailed balance for reversible binding interactions (unless rates are explicitly defined to violate this). Lastly, as for any spatial method, NERDSS must define boundary conditions for the simulation volume. While all NERDSS routines work within a simple rectangular volume and (if appropriate) flat membrane, the software also handles arbitrarily curved surfaces^35^. This new feature for single-particle RD^36^ also includes an efficient method for surface binding using an implicit lipid model. The implicit lipid model accurately reproduces binding kinetics from the more expensive explicit lipid simulations with orders of magnitude speed-ups^36^.

### NERDSS quantifies *in vitro* experiments of clathrin assembly on membranes

A powerful aspect of NERDSS is the ability to generate microscopic models where output can be directly compared with experiments of multi-component assembly. Recent *in vitro* experiments have measured the kinetics of clathrin localization to adaptor-protein coated membranes^29^. We initialize the simulations to match the experiment, with adaptor proteins bound to the membrane surface and a relatively dilute (80nM) but large volume of solution clathrin that is in excess of the membrane surface area (Fig 2). In our model, clathrin has three sites to bind other clathrin molecules, and three sites to bind the adaptor proteins, thus localizing the molecule to the 2D surface.

**Figure 2.**
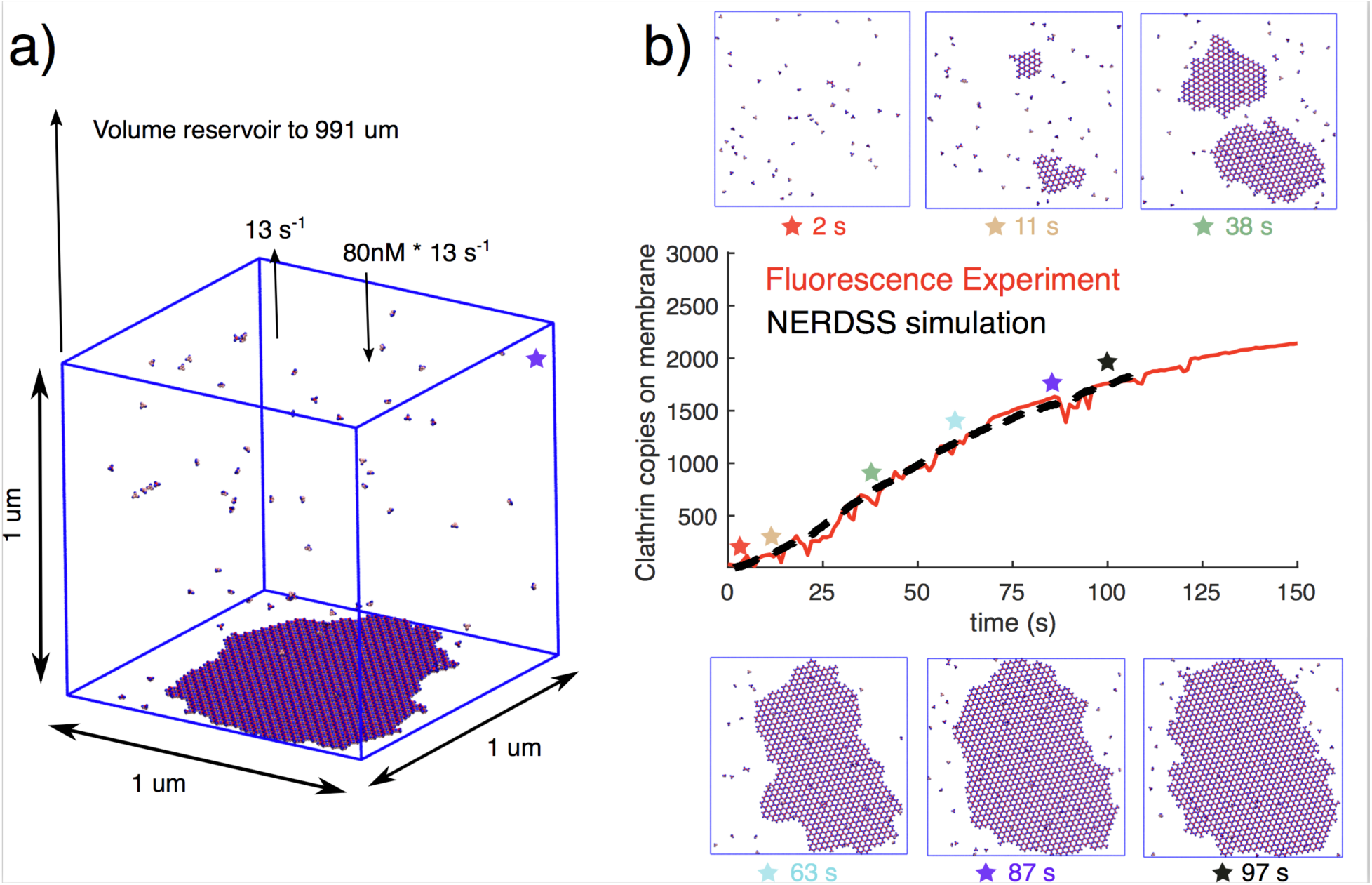
NERDSS simulations of clathrin assembly kinetics on membranes reproduce *in vitro* experiments. a) Simulations were set up to mimic experimental conditions from recently published work of Holkar et al^29^, where fluorescence of clathrin on membrane tubules was measured with time. A constant (fluctuating) solution concentration of clathrin (at 80nM) was maintained through exchange with the large volume reservoir. b) Fluorescence data averaged over multiple tubules is shown in red. The best model result is shown in black dashed. The y-axis reports clathrin copies on the membrane, per um^2^, where fluorescence data can be rescaled because it has arbitrary units. Despite the arbitrary fluorescence units, both the observed time-scales and the known geometry/concentrations of the experiment strongly constrained the models. Only a small subset of models could reproduce the slow lag (10s) in the experimental kinetics, followed by relatively rapid nucleation of the lattice on the surface, which slows due to a loss in accessible surface area and adaptor binding sites (Methods). Snapshots along the trajectory illustrate how a few distinct sites nucleate, which eventually grow radially outward to coat the surface. Each clathrin contains a center-of-mass and three ‘leg’ binding sites at 8 nm from the center, such that a bound pair has centers separated by 17nm. Three adaptor binding sites are offset from the flat clathrin interaction surface. No curvature effects are included here. Adaptor copies were initialized already bound to the surface at 0.007/nm^2^ (Methods).

The simplest model we created that can reproduce the observed experimental kinetics (Fig 2) has two sources of cooperativity. First, once clathrin binds the adaptor proteins, its affinity for other clathrins is increased, an experimentally well-established phenomenon^37^. Second, once clathrin is localized to the surface, it binds additional adaptors or clathrins with a 2D rate constant^18^. Both sources of cooperativity contribute to relatively rapid nucleation and stabilization on the surface, following an initial lag due to slow binding of clathrin to the adaptors. Saturation of clathrin on the surface results from a loss of accessible surface area and adaptor binding sites. The binding affinities we specify between the proteins are comparable to literature measurements (collected here^18^) (see Methods for full details). We use here flat clathrin trimers because the membrane is modeled as a flat surface that does not currently undergo any dynamics. Flat clathrin lattices do form *in vitro* and *in vivo*^38, 39^, supporting that at least some assembly is de-coupled from membrane remodeling^40^. Membrane bending could produce an additional source of cooperativity, if clathrins favor high curvature as well as inducing it, which is an active future direction for this software.

### NERDSS tests clathrin cage assembly designed by four distinct models of clathrin-clathrin interactions

Clathrin assembly in solution is observed experimentally either at low pH or in the presence of clathrin terminal binding adaptor proteins^37, 41^. We model distinct assembly conditions by changing binding strengths, cooperativity, and binding geometries of clathrin-only solutions, contrasting how multiple observables vary with distinct model parameterizations. In all systems, we include 100 clathrin triskelia, with each clathrin containing a center-of-mass and three ‘leg’ binding sites at 120 degree rotations from one another and at 10 nm from the center, such that a bound pair has centers separated by 21nm. The baseline model has a K_D_ for leg-leg binding of 100 *μ*M, which is comparable to experimental estimates of trimer dimerization^42^. Very little binding is observed in solution at this concentration (Fig 3a), which is consistent with *in vitro* studies of clathrin in solution at physiological pH^41^. The same results are observed in non-spatial simulations (Fig S3).

**Figure 3.**
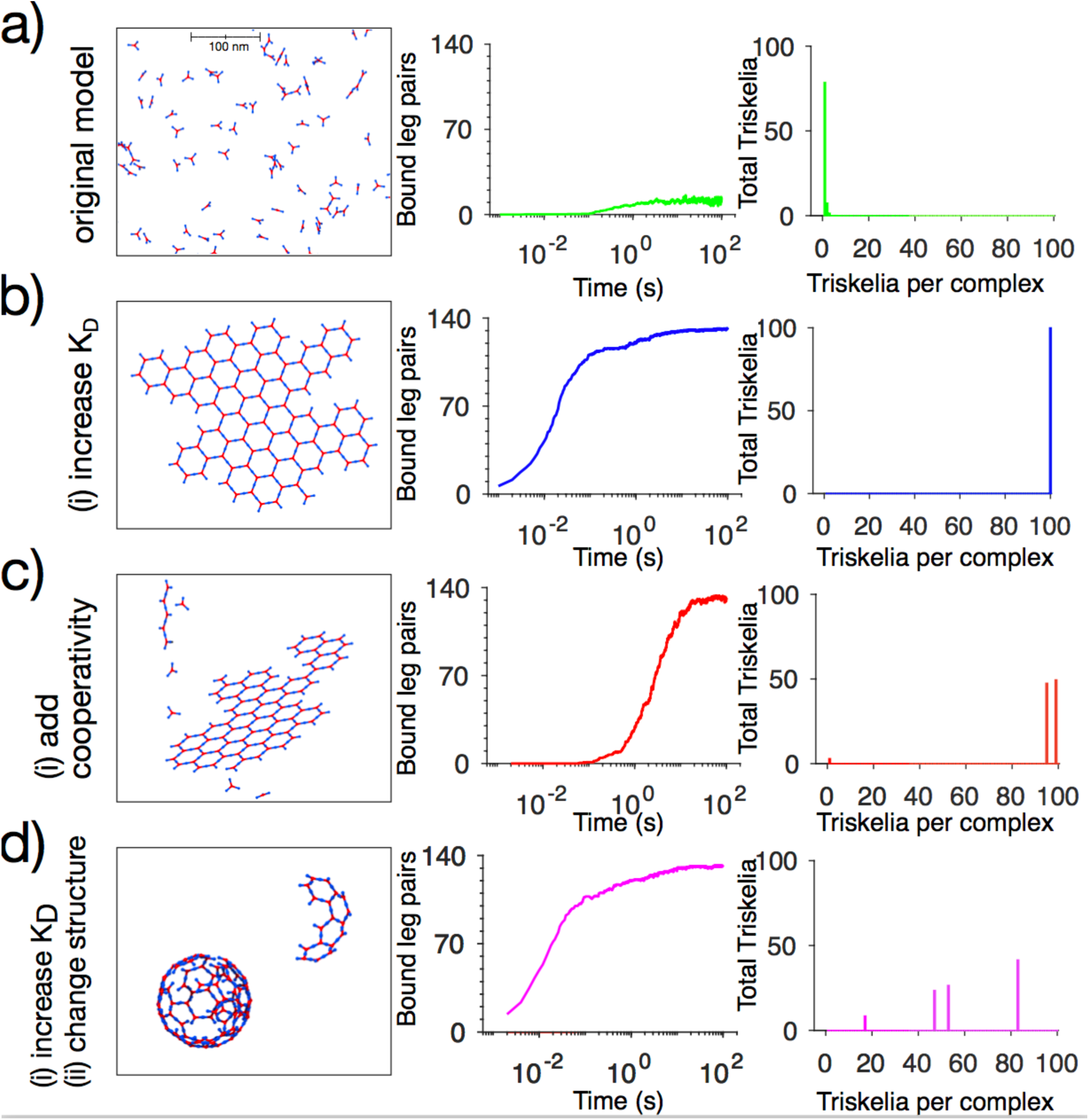
Simulations of clathrin-cage assembly in solution, with 1.3µM clathrin triskelia. a_1_) For K_D_=100µM, most clathrin remains monomeric in solution. a_2_) Counts of pairs of bound clathrin trimer legs. a_3_) Histogram showing the total copies of clathrin triskelia as distributed complexes of increasing size. All simulations have 100 trimers in them. b) By increasing the K_D_ to 0.2µM, large flat lattices form. c) By introducing cooperativity such that binding of monomeric to non-monomeric clathrin is stronger by a factor of 10, and non-monomeric to non-monomeric is again 10 times stronger, lattices can form despite a weak monomeric K_D_=100 µM. d) By changing the geometry of the clathrin monomers but keeping the same K_D_=0.2 µM from (b), they assemble into spherical cages with similar binding kinetics but that nucleate smaller complexes (d_3_ relative to b_3_). Results are averaged over 3-5 trajectories, and SI movies are generated with VMD.

By simply strengthening the K_D_ to 0.2 uM, flat clathrin lattices assemble quite rapidly (k_off_=1s^−1^), and by 100s they are in stable, giant components (Fig 3b). An additional binding parameter is allowed for all models that can result in loop closure (here, hexagonal loops). This parameter acts as a measure of positive or negative cooperativity due to the formation of two simultaneous interfaces within the closed loop, and determines the relative stability of closed vs open loops (SI Methods). This can thus alter the final equilibrium for a given dimerization K_D_, favoring fewer bonds and more remodeling (Fig S3-S4).

By instead introducing cooperativity in binding between clathrins, such that the rate depends on whether a monomer is already bound to another clathrin, flat lattices assemble after a delay, followed by rapid growth (Fig 3c). This model thus has three on-rates for clathrin-clathrin binding, with strongest binding between clathrins that are already bound to others. This growth model results in a broader distribution of lattice sizes (Fig 3c_3_). (Movie S1)

By taking the model of Fig 3b (K_D_=0.2 uM) and altering the geometry of a clathrin monomer to tilt legs 10 degrees below the plane, spherical cages form (Movie S2). The kinetics of leg-leg binding remains very similar to Fig 3b_2_, but the sizes of the complexes are smaller due to spatial constraints of closed spheres and steric overlap (Fig 3d_3_). Two important algorithmic features emerge in this model due to the rigid structure of the clathrin trimer and its assembly into a spheroid. First, because the lattice cannot form perfectly on the sphere, defects appear even within a single closed hexagon. Biomolecules typically have the flexibility to bend and form contacts by supporting a distribution of structural geometries, so we allow sites within a complex to bind one another even when they are not at perfect contact (Methods). These bonds are important because they stabilize the complex against dissociation, even though they do not change the rigid structure of the complex. Second, with defects present in the lattice, evaluating steric overlap upon association is trickier, as monomers may overlap but binding sites themselves are offset from one another (Methods).

### NERDSS probes lattice assembly driven by localization to the membrane

Increasing component concentrations is another natural mechanism to nucleate assembly. In Fig 4 we show how localization to the 2D membrane surface can, on its own, nucleate clathrin lattices by increasing the effective concentration of clathrin relative to the 3D solution (Movie S3). Similar to Fig 2, clathrin can be localized to the surface by an adaptor protein, but here the adaptor starts off in solution. The adaptor contains both a protein interaction motif for binding the clathrin terminal region, and a lipid binding domain that typically targets phosphatidylinositol bisphosphate, PI(4,5)P_2_, which is ∼1 mol % of PM lipids. The clathrin-clathrin interaction is set here to a weak value of 100*μ*M (Fig 3a), and thus minimal binding occurs in solution. Once the clathrin have been localized to the surface, however, they can assemble into a lattice that is stabilized by relatively weak clathrin-clathrin contacts but is nonetheless favorable due to the small search space available on the 2D surface. No cooperativity due to adaptor binding is included here.

**Figure 4.**
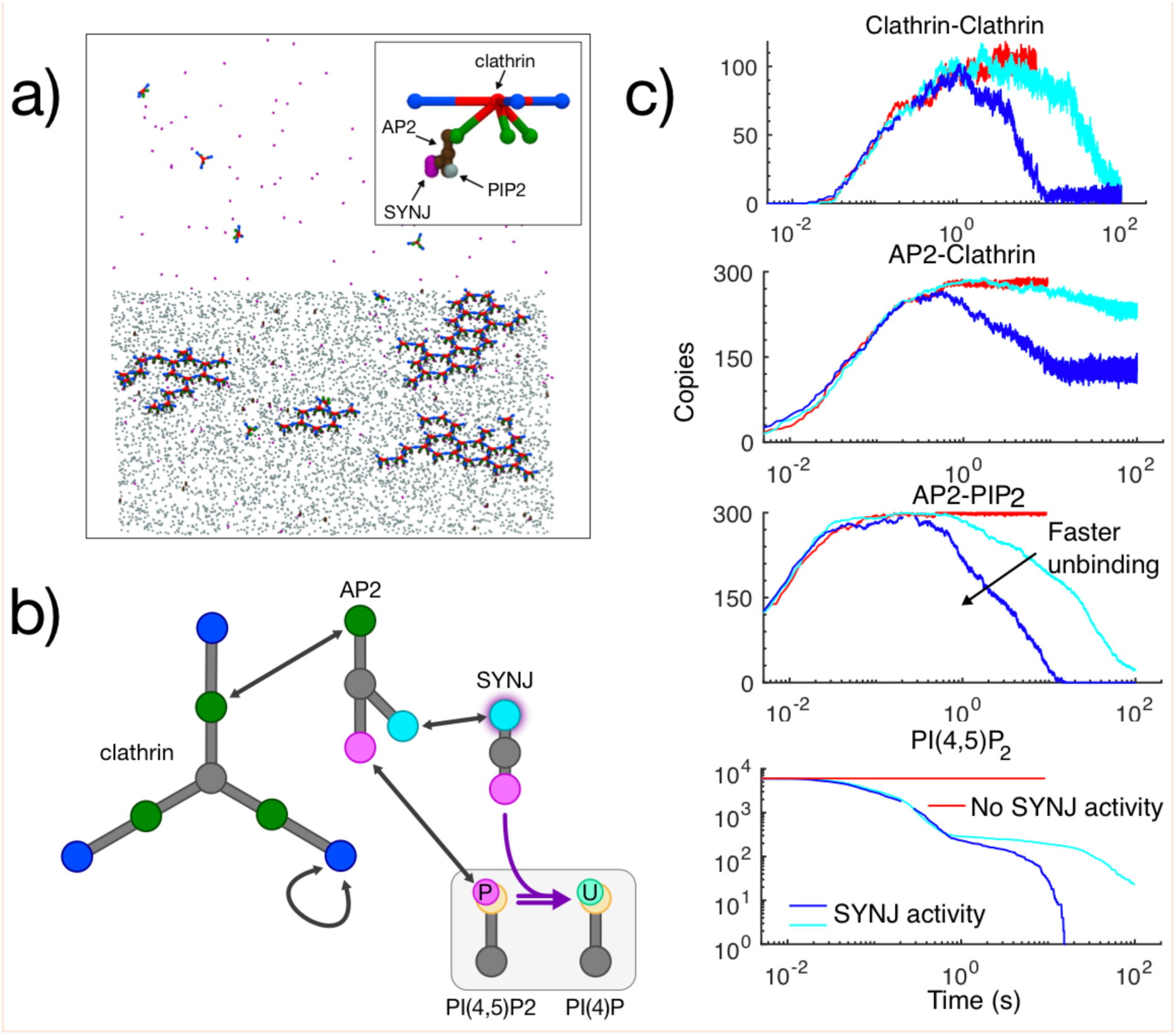
Clathrin assembly and disassembly by membrane localization and de-localization. a) Three multi-valent solution proteins are included in this model (clathrin, AP-2, synaptojanin) and one membrane lipid (PI(4,5)P_2_). For this model, clathrin does not assemble in solution even if bound to the AP-2 protein (similar to Fig 3a). Only upon localization to the 2D surface does clathrin become concentrated enough to form a lattice. b) Clathrin contains three sites for binding other clathrin, and three separate sites for binding to AP-2. The AP-2 protein has a dedicated site for each of the three other components, driving the clustering on the membrane. The synaptojanin phosphatase has a site for AP-2, and an independent site that targets the PI(4,5)P_2_. Critically, the AP-2 protein does not interact with PI(4)P, which is produced by the binding of synaptojanin to PI(4,5)P_2_. c) With no enzymatic activity, the assembly proceeds to an equilibrium steady state (red curves). With phosphatase activity turned on, the majority of the lipids are converted by 1 s, except the population that has been already bound to AP-2. The clathrin lattice thus has time to assemble, but is gradually destabilized as determined by the rate of unbinding between adaptor and lipid, and adaptor and clathrin, which increases from 1s^−1^ to 10 s^−1^ between cyan and blue.

### Enzymes can drive lattice disassembly by removing links to the membrane

By removing links between clathrin and the membrane, we can drive the clathrin assembly back into solution where our clathrin lattice is no longer stable. Physiologically, this is achievable by changing the phosphorylation state of the lipid PI(4,5)P_2_, which is essential for localizing adaptor proteins, and thus clathrin, to the membrane. Thus, without altering the clathrin or the adaptor proteins directly, we can drive disassembly of the lattice. We include here 10 copies of a lipid phosphatase (e.g. synaptojanin), which converts PI(4,5)P_2_ to PI(4)P irreversibly. The adaptor protein cannot bind to PI(4)P, thus removing its link to the surface. Importantly, the phosphatase synaptojanin can be localized to sites of clathrin-coated structures *in vivo* through protein interactions with the adaptor protein AP-2, thus allowing it to act in 2D to rapidly dephosphorylate unbound PI(4,5)P_2_. While *in vivo* it has been shown to play an important role in dephosphorylating PI(4,5)P_2_ after fission from the membrane^22^, we demonstrate here how it is capable of driving disassembly of plasma-membrane bound clathrin-coated lattices, at least in this *in vitro*-type model.

We find the timescales of the lattice disassembly are sensitive to the binding and unbinding kinetics of all protein-protein and protein-lipid interactions (Fig 4). In particular, although the phosphatases rapidly de-phosphorylate unbound PI(4,5)P_2_, the adaptor proteins protect the subset of lipids they are bound to, and their links are cut on a time-scale that is determined by unbinding of the adaptor-lipid interactions. Because clathrin can bind up to 3 adaptor proteins, its dissociation from the membrane then also depends on dissociation of all 3 adaptors from the surface. Although this model is not fully parameterized to reproduce experiment (as in Fig 2), it illustrates how localization, enzymatic activity, and binding kinetics can be tuned to control assembly and disassembly.

### Model building requires orientations for each pairwise binding reaction: application to viral assembly in solution

To illustrate the model design process, we consider here the Gag monomer, a retroviral protein that dimerizes with itself and oligomerizes into a hexagonal lattice on the plasma membrane of infected cells to form the so-called immature virion^30^. Gag can be coarsely approximated as a linear protein, which we specify via a homo-dimerization site H close to the center of mass, and two distinct oligomer sites G1 and G2 that define the molecule width, and that bind to each other. To form a lattice of hexamers from these monomers, two distinct types of dimers are sufficient, with relative orientations defined by five separate angles for each dimer (Fig 5).

**Figure 5.**
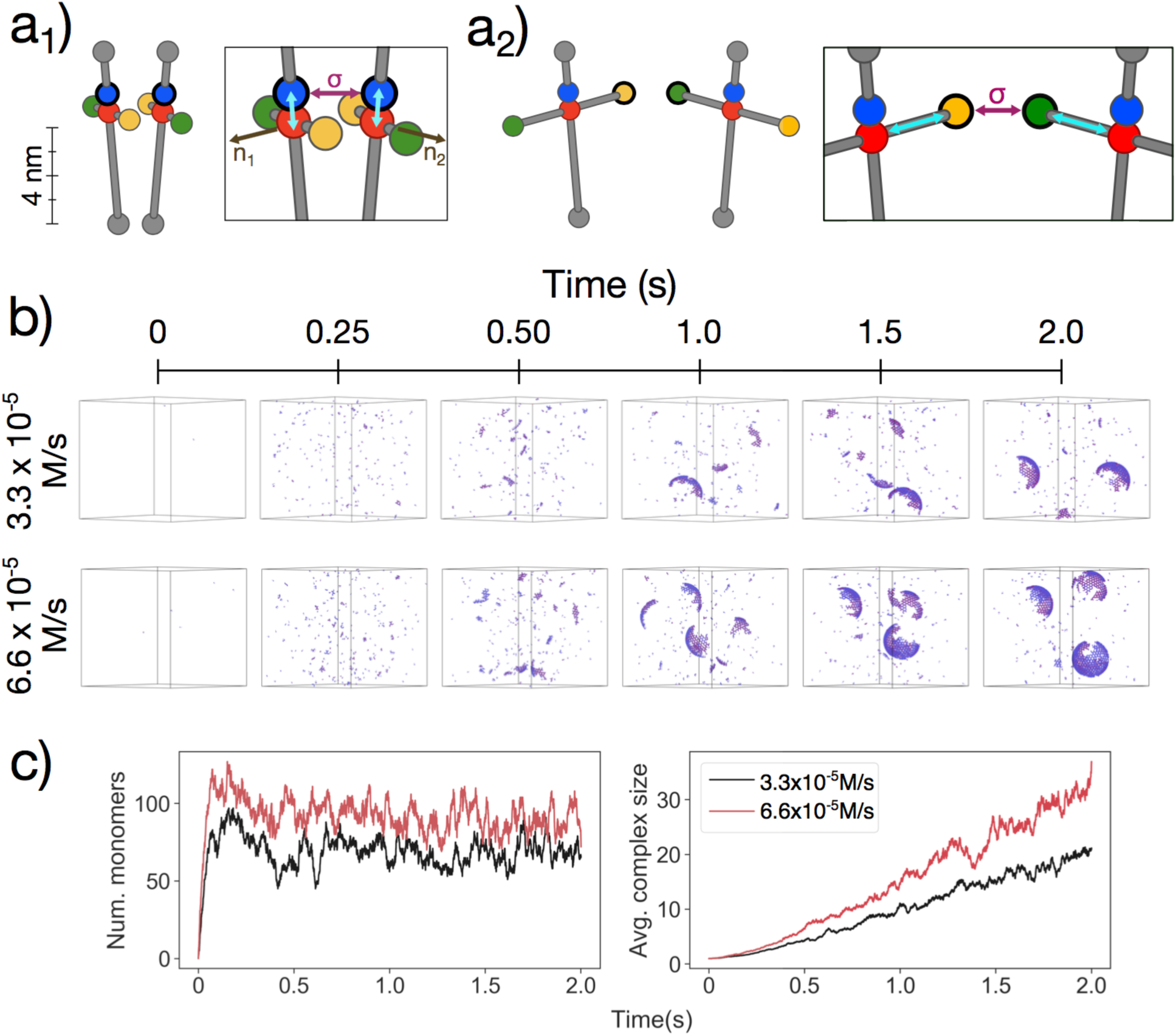
Gag monomer assembly model set-up and titration experiment. Each Gag protein monomer is given 3 active binding sites, and 2 inactive binding sites, positioned relative to a center-of-mass (COM) site in red. The blue site is for homo-dimerization, H, and the green (G1) and orange (G2) sites are for heterodimerization to produce hexamers. The positions of the sites are chosen to result in assembled structures with the proper separation between protein centers. The orientation of the bound homodimer is specified through 5 angles, and includes a slight tilt of one monomer relative to the other to ensure a spheroid lattice forms. The 5 angles are based on the COM to interface vectors (turquoise lines), the sigma vector (purple line), and two molecule normals (black vectors) that must not be co-linear with the interface vectors to provide an additional dimension. The orientation of the bound heterodimer similarly requires 5 angle definitions, and G1 must bind G2 at a cis orientation to ensure a hexamer loop can form (inset). b) Trajectories show the titration in of Gag monomers at 3.3 10^−5^ M/s and 6.6 10^−5^ M/s. c) The number of monomers initially grows rapidly, but reaches a relative steady-state due to assembly and to degradation, which occurs at a rate of 1s^−1^. Assembled proteins are protected from degradation, causing a continual growth in sizes of assembled complexes.

To illustrate the functionality of NERDSS, we initialize the volume with zero Gag monomers. We then titrate them in using a zeroth order reaction, where they are placed randomly in the simulation volume. Thus, the concentration slowly increases, favoring slower nucleation and growth of the Gag into the more stable homodimers, and eventually hexamers (Movie S4). Monomeric Gag is also degraded, although assembled Gag is not. Thus the long-time behavior consists of Gag as stable spheroids, or as short-lived (∼1s) monomers. Although Gag hexameric interactions are known to be auto-inhibited prior to binding to either RNA, negatively charged analogs, or the membrane^30^, we here treat all interactions as constitutive for simplicity. As designed, the Gag is able to assemble into large spheroids entirely due to the two specified reactions (Fig 5). The assembly kinetics depends on the titration speed, which can provide additional knobs to help optimize the model against experimental data.

### Recapitulation of protein expression oscillation for a circadian clock model using NERDSS

Beyond its ability to perform structurally-resolved simulations of self-assembly, NERDSS is also able to simulate other equilibrium and non-equilibrium phenomena. In Fig 6 we simulated a previously developed minimal model of a circadian genetic oscillator^31^ with NERDSS, as well as with PDEs and the stochastic simulation algorithm (via Virtual Cell software^1^ (SI)). The model recapitulates how the expression of two proteins, an activator protein (A) and repressor protein (R), can be coupled to produce robust oscillations of both proteins and their bound complex. For our NERDSS model, when mRNA and proteins are created, they are placed adjacent to the molecule creating them. The A-R bound complex is similarly represented via the points separated by their binding radius sigma, such that the complex is somewhat visible in the trajectory snapshots. All rates were accelerated by a factor of 3600 relative to the original rates, due to computational costs, such that oscillations occurred over seconds rather than hours time-scales. This change made the binding events diffusion limited, meaning they could become sensitive to the spatial distribution of particles. However, because the species mix rapidly and the volume is not too large (Movie S5), the kinetics were essentially insensitive to diffusion and the spatial dimension, agreeing quantitatively with both a PDE simulation and a non-spatial Gillespie simulation of the same model (Fig S3).

**Figure 6:**
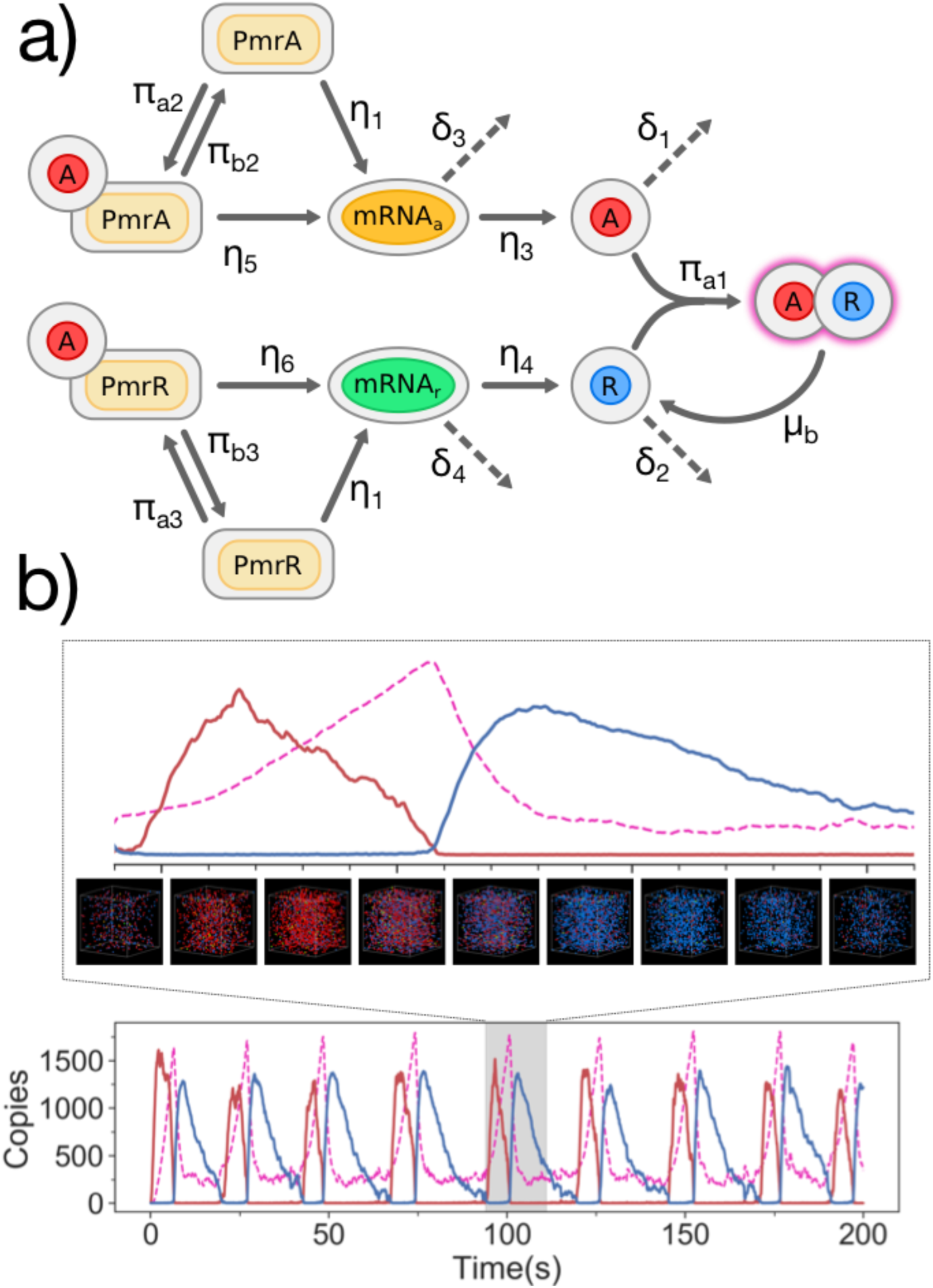
Spatially-resolved simulation of a circadian clock model illustrates oscillatory protein expression. a) The model contains 6 components, with the activator (A) and repressor (R) proteins produced by their corresponding mRNA, which are produced by a single copy of each gene (PmrA and PrmR). Unimolecular reactions (except dissociation) are shown with reactions indicated by *η* or *δ*. The binding and unbinding reactions are shown with rates *π*. b) Snapshots of simulations have particles colored according to part (a). Binding of R to A produces a complex A-R (magenta dashed), whose copy numbers oscillate in time, peaking between the A and R copy number oscillations. The A-R complex unbinds with a rate *μ*, that results in only R, as A is modeled as degraded instantaneously. The period of oscillations of A (and R) from NERDSS is in close agreement with non-spatial and PDE-based simulations of the same model, calculated as 24.5s, 24.8s, and 25.1s, respectively (Fig S5 and SI). Because of the constant production and degradation of components in this model, we used Ovito software to produce movies for this model (Movie S5) and the Gag model (Movie S4).

## Discussion

NERDSS has several advantageous features which make it a powerful and immediately useful tool for cell-scale simulations. First, NERDSS is transferrable between distinct systems thanks to its rate-based interaction framework, which avoids the time-consuming parameterization of energy functions often hard-coded for specific assembly systems^11^. Although protein geometries and orientations of bound complexes in NERDSS are system specific, we provide a GUI to facilitate user-design of proteins and their bound states. Second, unlike existing spatial rate-based approaches^13^, the built in molecular structure of NERDSS not only enforces excluded volume of binding sites, but evaluates whether steric overlap or volume restrictions would prevent the formation of unphysical assembly structures. Third, it uses the Free-Propagator Reweighting (FPR) algorithms^12, 16, 17^ to efficiently propagate species while retaining accurate rates of association and dissociation, allowing for current simulations of timescales on the order of seconds to minutes. NERDSS converts carefully between microscopic rates used in FPR and macroscopic rates commonly defined from experiment, with extensive validation^16^. Fourth, NERDSS uses a BioNetGen Language (BNGL)-style syntax^43^, and models built in other software packages using similar syntax can be ported to NERDSS with minimal alterations, making it available for immediate use. In this combination structural resolution, efficient propagation and accurate treatment of association rates, user-friendly input file syntax, and graphical user interface (GUI), NERDSS is a unique tool for simulating equilibrium and non-equilibrium self-assembly and other cell-scale phenomena.

NERDSS is designed to be extensible to not only users but to developers expanding functionality. NERDSS is built off of algorithms that support, for example, the implementation of more elaborate physical models that introduce electrostatic interactions between particles^12^. A current limitation of NERDSS is that all species must be rigid bodies, and thus assembly defects within a complex are accommodated approximately by using a distance tolerance. Molecule flexibility is a natural extension beyond rigid molecules that would improve defects and also enable better treatment of disordered regions and genomes. A particularly exciting future development is integration of NERDSS with continuum membrane models^36^, to allow more realistic simulations of vesicle and virion formation dynamics and coupling of assembly to mechanical force generation. Ultimately, NERDSS has encoded as a core feature the ability to resolve relatively fast processes over long time scales, and individual proteins over long length scales, simulating otherwise intractable protein assembly dynamics. This provides immediate use in helping overcome the challenges of understanding or designing self-assembling structures in biology.

## Methods

### Implementation

NERDSS is written in ANSI/ISO standard C++11 and is available on Linux and macOS. The accompanying GUI is implemented in Java. The source code for both can be found at https://github.com/mjohn218/NERDSS and is provided under the GNU general public license (GPL). A user guide and simulation example systems can be found on the software page. Input files are formatted similar to BNGL style, such as are used in RuleBender^44^ and the NFSim software^3^ and is compatible with Virtual cell^1^. Additional features are necessary, however, due to the spatial and structural details used in NERDSS.

### Reactions

NERDSS supports the three primary types of reactions: zeroth-, first-, and second-order. The rates of these reactions, other than creation, can depend on the states and/or pre-existing interactions of any participant species’ interfaces.

### Zeroth-order Reactions

*De novo* particle creation reactions are treated as a Poisson process. No events occur if a *URN* < exp(−*λ*), where *λ*= *k*_0_VΔt, *k*_0_ is the rate in units of M/s, V is the simulation volume (in units of M^−1^), and Δt is the timestep. Therefore, *N* events occur based on 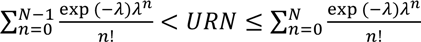. Created particles are placed within the simulation volume with random coordinates. Zeroth-order reactions can be used to titrate species into the simulation volume.

#### First-order

First-order reactions are treated as Poisson processes, with a reaction probability per each interface of *p*_1_(Δ*t*) = 1 − exp(−*λ*), where *λ*= *k*_1_Δt, *k*_1_ is the microscopic rate in units of s^−1^ and Δt is the timestep. The reaction rates can be specified either as microscopic rates or as macroscopic rates. Only one such event can occur per timestep per molecule. Four types of first-order reactions are supported:

1. Creation: Create a copy of a particle B from another particle A, where interfaces required such as (a) are indicated within the parentheses: Examples are transcription or translation, e.g. A(a)→A(a)+B(b)
2. Destruction: Destroy a molecule and all other molecules in its complex. If the interfaces (a) are specified as unbound, it will only destroy unbound molecules. E.g. A(a)→NULL
3. State change: Change a state of an interface on a molecule, e.g. A(a∼p)→A(a∼u)
4. Dissociation: Remove an interaction between the interfaces of two particles, written as the conjugate back reaction of an association reaction. To retain detailed balance, the particles are left such that the dissociating interfaces are at their binding radius, e.g. A(a!1).B(b!1)→A(a)+B(b)

#### Second-order Reactions

Bimolecular reactions are applied to specific interfaces between two proteins. Using BGNL syntax, association events can be conditional on the state of the interface, e.g. occurring only when the binding interface is phosphorylated, or only if the protein is bound through another interface as well.

As FPR reaction probabilities are invariant with respect to orientation, the reacting particles are “snapped” into place in a predefined geometry in order to prevent arbitrary structures from forming, with interfaces always placed at the binding radius σ from one another. The geometry is defined by a set of vectors and five angles between and within each particle. If angles are undefined, interfaces bind to a separation of σ at the orientation they were when the event occurred. Both molecules are translated and rotated into place based on their relative translational and rotational diffusion constants. Thus, smaller complexes will move more to orient, and complexes restricted to the membrane do not rotate out of their membrane-localized orientation. Detailed definitions of these vectors and angles can be found in the Supporting Information.

Reaction rates can be defined either as macroscopic rates, in units of nm^3^/s, or as microscopic or intrinsic rates, in units of nm^3^/s (converts to M^−1^s^−1^ with multiplication by 0.602). For reactions in 2D, the software defines the 2D rate for two interfaces based on their 3D rate divided by a lengthscale, which by default is set to 2σ (units of nm), but which can be independently specified per reaction using an input parameter. A reaction is identified as 2D when it involves two species that have no diffusion in *z* (e.g. lipids), or between interfaces on two complexes that are localized to 2D (D_z_=0). For self-binding reactions, and for reactions where one reactant is in 2D, we correct for factors of two between macroscopic and microscopic rates. A Table in SI explains these relationships. Probabilities of each reaction are calculated with the FPR method as previously described, and are parameterized by the intrinsic reaction rate *k*_a_, the net diffusion coefficient of the reactants *D*_tot_, and the binding radius σ. Two types of bimolecular reactions are supported:

a. Association: Form an interaction between two interfaces of two molecules, which, if the association is reversible, has a conjugate first-order dissociation reaction. The resulting complex is then treated as one unit for future propagation. E.g. A(a)+B(b) → A(a!1).B(b!1)
b. State change: Change the state of an interface on a particle, facilitated by another particle, such as phosphorylation by a kinase. This is thus a binding reaction that automatically results in a first-order state change of one interface, and both reactant species remain unbound. E.g. A(a)+B(b∼u) →A(a)+B(b∼p).

### Intra-complex binding

An important consequence of forming self-assemblies is that molecules can be capable of binding to one another through free interfaces when they are within the same complex. These intra-complex binding events involve two interfaces becoming a bound complex, but are not truly bimolecular, as they do not involve a search to find one another—they are already co-localized. They are thus unimolecular, or first order reactions, and cannot be treated with the same binding probabilities as is used for bimolecular events. We define the binding probability of these intra-complex or loop closure events thus using a Poisson probability, similar to the 1^st^ order reactions described above. Then we must specify a unimolecular rate, given a bimolecular rate constant. We define this rate such that the equilibrium between the bound and unbound states for a two-step process of loop closure, that is, a bimolecular event and a unimolecular loop closure, is the same as if the protein closed the loop in a single step, forming both bonds at once. The rate is given by:

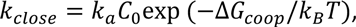

where *C*_0_ is the standard state concentration of 1M. For loop closure the user can thus specify one additional parameter, Δ*G_coop_*. Positive values of this enable more dynamic reversible loops, which otherwise tend to be highly stabilized relative to single bonds. We derive this expression in the SI.

We allow these events to form when the interfaces are spatially proximous, with the user able to specify a maximum distance to allow these events to occur. The maximum distance is the binding radius, σ, multiplied by a factor, bindRadSameCom, which by default is set to 1.01, which includes only perfectly (with numerical precision errors allowed) aligned contacts. The primary reason to increase it is because for some self-assembly structures (such as the curved clathrin cage), the rigid structures cannot form perfect, defect-less structures. Allowing binding between legs that are close together mimics structural flexibility present in the biological molecules. Finally, the software prohibits binding between a pair of molecules if they are already bound through a distinct set of interfaces. Hence, intra-complex or loop-closure events must be mediated through at least a third molecule (e.g. 4 molecules in the case of clathrin lattices).

### Evaluation of Steric Overlap

Once association events occur between two interfaces, their two parent complexes are rotated into place to generate the proper, pre-defined geometry of the two binding proteins. The exception is the intra-complex binding, where because the binding happens within a single complex, no rotation or movement occurs. For two separate complexes, however, this rotation into place can result in steric overlap between interfaces that are part of the complex but not part of the binding event. We evaluate overlap after association by measuring distances between all centers of mass in the new complex, and defining a minimum distance, overlapSepLimit, that will determine steric overlap. If any pair of proteins have COMs less than overlapSepLimit, which by default is 1nm, the binding event is rejected, and the two complexes instead undergo a diffusion move during the time-step.

A final steric overlap check is then performed between the newly bound complex, and all other binding interfaces of complexes present in the simulation, to ensure excluded volume is maintained. If the new complex produces steric clashes with other binding interfaces in the simulation volume, the binding event is rejected.

### Boundary effects

By default, the simulation boundaries are reflecting. If association events result in a very large lattice, it is possible for the structure to extend beyond the physical boundaries of the simulation volume. These moves are thus rejected. On a related note, if association events result in very large rotational re-orientation of a complex, these moves can also be rejected, as they result in an unphysical amount of displacement per time-step.

### Diffusion constants of complexes

Once proteins bind to form a new complex, their translational and rotational diffusion constants are updated to reflect the larger hydrodynamic radius of the bound complex. Each protein molecule has a user defined D_t_ and D_R_. Once a bound complex forms, the new transport coefficients are then defined based on all *N* components of the complex by simply assuming the radii sum:

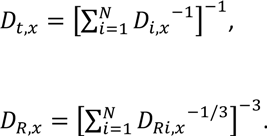

and the same in y and z. For molecules restricted to the surface D_t,z_=0 and all rotational diffusion except (if desired) D_R,z_ are also zero. Correspondingly, any complex that contains these molecules will also have that diffusional component set at zero.

### Data output

NERDSS produces restart files which ensure that when simulations are interrupted for any reason, they can be restarted exactly from where they left off, similar to Molecular Dynamics software. NERDSS also records several system properties in time, including the coordinates of all species, and the distribution of proteins amongst different types of complexes.

### Simulation algorithm steps

*(i)* <u>Initialization.</u> Copies of seed particles are created from the provided templates, given random coordinates, and checked for placement within the simulation volume and overlap with other particles. Overlap is only corrected for if two interfaces which can react with one another are within the binding radius of their reaction. If two particles are found to be overlapping, one of the particles is reinitialized with new random coordinates and all particles are rechecked for overlap.
*(ii)* <u>Optimization.</u> To optimize evaluation of the two-body (bimolecular) events, the simulation volume is split into *N* sub-boxes, where the length of each sub-box edge is at least 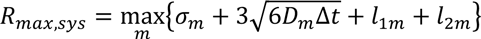, where *m* loops over all binding reactions, σ is the binding radius, *D* the total diffusion coefficient of both reactants, and *l*_im_ is the radius of the reactant protein *i*. Molecules are assigned to boxes based on the position of their COM. This allows for looping over all possible binding partners only in a molecule’s current and neighboring boxes, without excluding any partners with a non-zero reaction probability.
*(iii)* <u>Reactions.</u> Within each time-step, reaction probabilities are calculated in order from zeroth-, first-, and second-order reactions.

For zeroth-order reactions, a number of molecules *N* to create during that step is determined, and then these new molecules are placed in the simulation volume randomly, ensuring that they do not overlap with other binding partners.

For first-order reactions, each molecule is checked against the list of reaction rules for compatibility, i.e. if its interfaces are in the correct state and if it has the correct bound partners, and the probability checked against a uniform random number. If a reaction occurs, it is performed immediately and the molecule and its interfaces cannot take place in any further reactions or diffusion during that time-step.

For second-order reactions, pairwise reaction probabilities are calculated for each pair of interfaces within R_max_ of each other that are free to bind. Next, for each molecule, probabilities for each of their possible binding reactions during that step are then checked against a uniform random number. Should a reaction occur, it is performed immediately. For all reactions, if a molecule is created or involved in a reaction in any way, including as a non-reacting member of a reacting complex, it cannot participate in any other reaction during the course of the time-step, nor can it diffuse. However, other diffusing particles will include them in overlap checks. On the other hand, if a particle is determined to not be involved in a reaction, Gaussian distributed rotational and translational propagation vectors are chosen and stored for that complex.

*(iv)* <u>Propagation and overlap</u>. After all reaction checks are completed, interfaces which were within R_max_ of a reaction-partner during the time-step are checked for overlap. Since a multi-component complex only moves as a rigid unit, each complex with reactive interfaces within is looped over, along with a stored list of all reaction partners. If the newly displaced positions overlap (with displacements sampled from Gaussians for translational and rotational diffusion for both reaction partners), the displacements of complex *i* and its partner are re-sampled, and the overlap check restarts. Once a complex’s position has been updated, it cannot be moved again. For highly crowded simulations (not studied here) this pairwise sequential evaluation of overlap may not prevent all overlaps. We thus have another version of overlap evaluation that finds all reaction partners of a complex *i*, and all reaction partners of those partners, etc, to create a cluster of interfaces whose positions must be simultaneously solved to prevent overlap. This cluster overlap version can be slower and if a solution is not found in a few iterations, it proceeds to update positions by taking repeated smaller steps until the full time-step is reached.

### Simulation Details

All simulations were run on Dell workstations running Linux, with Intel cores (i7-8700, Xeon E5-2697v4, i9-9980XE), or MacBook Pros with Intel Core i7 7700HQ, or the MARCC supercomputer at Johns Hopkins University.

#### Simulations of Holkar *et al*. fluorescence experiments

Experimental data was kindly provided by Prof. Pucadyil. In Figure 2 we show the average over 40 kinetic traces of clathrin fluorescence accumulating on membranes pre-equilibrated with AP-2 *β* appendages. The experimental data could be fit, as in the original publication^29^, to an exponential with a lag time: *y*(*t*) = *b* + *H*(*t* − *τ*)*A*[1 − exp (−*k*(*t* − *τ*))], where *H* is the Heaviside function, *τ* is the lag time and *k* is the growth rate. The offset *b* and plateau *A* are in arbitrary units. For the averaged data, we found *τ*=10.7s and a k^−1^=107s. Similar results are found if all 40 curves are independently fit and then parameters are averaged. Simulation data was fit to the same function, where the best simulated model had *τ*=11s and a k^−1^=97s. The other parameters were *b*=54 and *A*=2823 (copy numbers/um^2^). The experimental data was plotted on the same y-axis (i.e. in units of copy numbers, not arbitrary units) by zero-ing the offset (subtracted off 290), and rescaling the height by 1.6.

The experimental V is 200*μ*L and the experimental A of 2.017×10^8^ *μ*m^2^ is calculated based on 1 nmol of total lipid, with an average lipid SA of 0.67nm^2^ and a two-leaflet bilayer formed. The experimental V/A ratio is thus 991 *μ*m. The NERDSS simulations were performed in a box that needs to maintain the same V/A ratio. We defined a box with a flat surface area (SA) of 1 *μ*m^2^, which would require a height of 991 *μ*m. The 80nM of clathrin in this volume thus equates to 47760 copies. The clathrin is in great excess of the available surface area; assuming each clathrin trimer takes up ∼201nm^2^ of space on the surface, maximally ∼5000 can be accommodated on the surface. To avoid propagating the thousands of solution clathrin, we instead set up the simulation with a height of 1 *μ*m, and maintained a constant (stochastically fluctuating) concentration of [Cla]_tot_=80nM (48 copies) in solution. This thus assumes that the total concentration of clathrin in the solution does not change as clathrin accumulates on the membrane, which is true to a very good approximation (total solution copies would drop from 80nM to 77nM over 100s). This was done by having a zeroth order reaction produce clathrin at *k*_create_=[Cla]_tot_*D*_Cla /_z^2^, to approximate the time-scales of diffusive flux across volume elements of height z, and by degrading the solution clathrin at a rate of *k*_destroy_=*D*_Cla_ /z^2^, where *D*_Cla_ is set to 13 *μ*m^2^/s. Clathrin on the membrane is not affected by the degradation reaction. *D*_R_=0.03 rad^2^/s. A time-step of 3 *μ*s was used.

Of the lipids on the surface, only 5 mol % were Nickel-chelator lipids (0.0746/nm^2^) that could bind His-tagged adaptor proteins. However, the affinity between the His-tag and the chelator lipid is weak enough^45^ that we expect only a fraction of these chelator lipids bind to adaptors (initially present at 200nM). For a K_D_ ranging from 1-10*μ*M between chelator and His-tag, we would have ∼1400-12000 adaptors on the surface before the clathrin is flowed in. We thus treat this as a parameter in the model optimization, with the models that best describe experiment containing between 6000-9000 adaptors on the surface. The model in Fig 2 has N=7000 adaptors affixed to the surface.

These simulations were initialized with zero clathrin, but the creation resulted in a steady-state concentration of 80nM by ∼0.2s. The ∼10s lag is thus not due to diffusion to the surface, but by slow binding of the clathrin to the membrane bound adaptors. This is consistent with the experimental findings^29^ that a different adaptor nucleated clathrin on the membrane with a much shorter lag time of ∼1s, presumably owing to faster binding of clathrin to this adaptor protein.

The clathrin-clathrin interaction strength was set to 120 *μ*M^42^, with an off rate of 10 s^−1^. Cooperativity was introduced for clathrin-clathrin interactions based on whether clathrin was bound to adaptor; if a trimer bound to an adaptor, its binding affinity for other clathrin trimers was increased across its other clathrin binding sites, by increasing the on-rate by a factor of 15, with the off-rate kept the same. The binding between clathrin and adaptor protein was constant and set to 25 *μ*M, similar to biochemical measurements^46^, with a slow on-rate of 4000 M^−1^s^−1^.

Simulations were run by treating the adaptor binding sites on the surface using our implicit-lipid model to speed-up the simulations^36^. This model has been shown to accurately reproduce the kinetics of binding to explicit surface sites in 3D and 2D^36^. Importantly, it captures 2D binding interactions between clathrin that are already localized to the surface via one adaptor, but have 2 additional binding sites available to bind further adaptors. All 2D binding rates were defined based on their corresponding 3D rates, with a length conversion factor of 30 nm^18^. This lengthscale applies to both clathrin-adaptor interactions in 2D, and clathrin-clathrin interactions in 2D. The clathrin loop cooperativity factor was set to *f*=0.001 (*f*=exp[-ΔG_coop_/k_B_T]).

#### Designed clathrin assembly simulations in solution

For Figure 3, all simulations were performed with 100 clathrin trimers in a box of length 0.494 *μ*m per side. The off-rates for all systems were set to 1 s^−1^ for all interactions. The K_D_ are 100*μ*M or 0.2*μ*M, with corresponding on-rates. The time-step was 0.2 *μ*s, and in Fig S3 we verify the same kinetics with a smaller time-step. The loop cooperativity factor was set to *f*=0.001, which makes the binding free energy about 1.5 times stronger than a single binding event, allowing more reversible loop formation events. In Fig S4 we show how this influences the kinetics and equilibrium. For the cooperative simulations, the on-rate between two clathrins is increased by a factor of 10 if one of the clathrin is not a monomer, and then increased by an additional factor of 10 if both clathrin are non-monomeric. Flat clathrin molecules are here designed to have a leg length of 10 nm, such that the distance between two centers of a bound dimer is 21 nm (includes the binding radius σ=1nm). The diffusion constants of clathrin were defined based on its size as D_t_=13 *μ*m^2^/s and D_R_=0.03 rad^2^/s, assumed isotropic. For puckered clathrin, the leg length was set to 7.5 nm, and the legs were offset from the plane by 10 degrees. For the puckered clathrin simulations, because defects could form on the lattice, we allowed binding to form (e.g. to close a hexagon), when the binding sites were within a distance of 5nm. Perfect contact occurs only in the flat lattice, where all clathrin trimers align to contact at the binding radius, σ=1nm.

The time-courses and complex histograms are averaged over 3-5 independent trajectories. The histograms shown are taken from the last steps in the simulations, although they are printed throughout.

#### Lattice assembly and disassembly on membrane simulations

Simulations from Figure 4 included 100 clathrin trimers, 300 adaptor proteins, 6000 initial copies of PI(4,5)P_2_ on the surface, and 10 copies of the phosphatase Synaptojanin in a box of size [0.7, 0.7, 0.494] *μ*m. The time-step was 0.2 *μ*s. The equilibrium constants for binding were consistent with literature observations, but they were not fixed by the model describing the experiment in Fig 2. Clathrin-clathrin interactions were set at 110*μ*M, with a 10s^−1^ off-rate in all models. No adaptor-driven cooperativity was included to isolate the role of 2D localization in driving assembly^18^. Clathrin-adaptor interactions had a relatively fast on rate of 6×10^6^ M^−1^s^−1^, and either a 1s^−1^ or a 10s^−1^ off-rate. The adaptor binding to PI(4,5)P_2_ also had an on-rate of 6×10^6^ M^−1^s^−1^ and either a 1s^−1^ or 10s^−1^ off-rate. The binding of the Synaptojanin to the adaptor protein was 6×10^5^ M^−1^s^−1^ and a 1s^−1^ off-rate in all simulations. Finally, the Synaptojanin bound to the PI(4,5)P_2_ at a rate of 2.5×10^7^ M^−1^s^−1^, except in the “No activity” simulations, where it was set to zero. Binding immediately results in dephosphorylation of the PI(4,5)P_2_ and freeing of the Synaptojanin to act again on another lipid. The lipid PI(4,5)P_2_ only has a single binding site, meaning that if it is bound to the adaptor protein, it cannot be acted on by Synaptojanin. The adaptor proteins cannot bind to the dephosphorylated lipid, PI(4)P.

Diffusion constants are set to *D*_cla_=13 *μ*m^2^/s, *D*_R,cla_=0.03 rad^2^/s. *D*_ap_=25 *μ*m^2^/s, *D*_R,ap_=0.5 rad^2^/s. *D*_pip_=1 *μ*m^2^/s (*D*_z,pip_=0), *D*_R,pip_=0. *D*_syn_=25 *μ*m^2^/s, *D*_R,syn_=0.5 rad^2^/s.

For all 2D binding interactions, where rates and K_D_s have units of per area (rather than per volume), we assigned rates based on dividing k_a,3D_ (the reaction on-rate) by 2σ and leaving k_b_ (the off-rate) unchanged. The loop cooperativity factor (applies only to clathrin-clathrin interactions) was set to *f*=0.001.

### Gag monomer assembly simulations

Gag simulations were run in a box of size [1, 1, 1] *μ*m. A time-step of 0.1 *μ*s was used. The homo-dimerization rate was set to 3×10^6^ M^−1^s^−1^ with an off-rate of 1s^−1^. The hetero-dimerization rate (G1+G2) is 10 times slower, at 3×10^5^ M^−1^s^−1^, with the same off-rate 1s^−1^. The transport coefficients were set to *D*_t_=10 *μ*m^2^/s and *D*_R_=0.05 rad^2^/s. Gag monomers are titrated into the system at a rate of 3.3×10^5^ M^−1^s^−1^, or twice as fast. Monomers (only fully unbound Gag), are degraded at a rate of 1s^−1^. Hence Gag that has formed a dimer or higher order complex is protected from degradation. The loop cooperativity factor was set to *f*=0.001. Gag proteins that were within the same complex could bind one another (e.g. to close a hexamer) even if they were not perfectly aligned, within a distance cutoff of 1.5nm (perfect contact is at the binding radius σ=1.0nm).

### Clock oscillator model simulations

The circadian clock model simulated here is taken directly from a published 2002 model^31^ that has been widely studied since then. We note here the changes performed for NERDSS. The units of all rates were changed from hr^−1^ to s^−1^, because resolving all collisions over hours-long time-scale is too costly. All 6 species in the system are point particles with a diffusion constant of *D*_t_=10 *μ*m^2^/s. Simulations were initialized with a single copy each of PrmR and PrmA. All creation and degradation reactions had the same values as published. Because the bimolecular association events were diffusion-limited between A and both genes (PrmR and PrmA) (6.02 x10^8^ M^−1^s^−1^) and between A and R (1.204 x10^9^ M^−1^s^−1^), the binding radius σ for these reactions had a minimum size >1nm, indicating an effectively longer length-scale over which these molecules can find one another. For A+PmrR, we set σ=5nm, and for A+R we set σ=8nm. This radius also sets the excluded volume of each of these pairs of molecules relative to one another. When RNA or protein are created, they are placed at the separation σ=5nm from their creator molecule, to minimize potential overlap in subsequent creation events. A time-step of 10 or 50 *μ*s was used, giving the same result.

For reversible bimolecular association events in NERDSS, we must define the intrinsic rates *k*_a_ and *k*_b_ such that^12^ 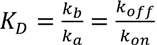, where the macroscopic rates *k*_PB_ and *k*_Pvv_ are the values defined in the model above and used in the PDE and the non-spatial stochastic simulation algorithm. One exception is the A+R→A.R event, which is not reversible. The intrinsic rate *k*_a_ is still defined in the standard way given the macroscopic on-rate^12^. The ‘unbinding’ event is actually a degradation reaction, where A.R→R, as A is degraded. The rate for this reaction thus is specified using the macroscopic rate of 1 s^−1^, not an intrinsic rate k_b_.

## Acknowledgements

MEJ gratefully acknowledges funding from an NSF CAREER Award #1753174 and an NIH MIRA Award R35GM133644. We would like to thank Danny Evans for setting up the Gag model, Ipsita Saha for providing feedback on Gag simulations, and Nomongo Dorjsuren for formatting the output files from the GUI. We thank Prof Thomas Pucadyil for very helpful discussions on the clathrin fluorescence experiments.

